# Intranasal Dantrolene Nanoparticles for Treatment of Amyotrophic Lateral Sclerosis as a Disease-Modifying Drug

**DOI:** 10.1101/2025.05.21.655232

**Authors:** Piplu Bhuiyan, Yutong Yi, Betty Wei, Allen Yan, Lola Dong, Huafeng Wei

**Author notes:** Correspondence should be addressed to Huafeng Wei.

## Abstract

Calcium dysregulation, caused by pathological activation of ryanodine receptors, contributes to motor neuron degeneration, motor dysfunction, and muscle weakness in SOD1-G93A transgenic amyotrophic lateral sclerosis (ALS) mice. This study investigates the therapeutic efficacy of intranasally administered dantrolene nanoparticles, a ryanodine receptor antagonist, on motor neuron function, muscle strength, spinal cord degeneration, and survival outcomes. Male and female C57BL/6SJLF1 non-transgenic control and SOD1-G93A ALS transgenic mice were assigned to one of three experimental groups: 1) NO TX: No treatment control; 2) IN-DAN: Intranasal administration of dantrolene in the Ryanodex formulation vehicle (RFV), at a dosage of 5mg/kg, administered daily from ages 90-120 days; 3) IN-VEH: Intranasal administration of RFV alone (as a vehicle control), following the same dosing schedule as the IN-DAN condition. Body weight and general motor function were monitored weekly, with survival recorded daily throughout the treatment period. At the treatment conclusion, neurological function was comprehensively evaluated using a standardized neurological scoring system. Motor coordination and balance were assessed using the balance beam test (beam widths of 12 mm and 6 mm) and the rotarod test. Muscle strength was evaluated by measuring grip force using the Kondziela inverted screen test. After behavioral testing, spinal cord tissues were collected for analysis. The levels of neurofilament light chain (NFL), a skeletal neuron protein, and spinal cord weight were determined to measure spinal cord degeneration. Compared to non-transgenic control mice, SOD1-G93A mice exhibited significantly elevated neurological scores, indicating severely impaired neurological function. This deterioration was robustly and significantly ameliorated by IN-DAN treatment by 90% (P<0.0001). Similarly, ALS mice demonstrated impairments in motor coordination and balance on the beam balance test, with dramatic and significant increases in crossing time and the number of foot slips. These impairments were greatly and significantly mitigated by IN-DAN treatment, by 78% in crossing time (P<0.0001) and 84% in the number of slips (P<0.0001) on the 12 mm-wide beam, but not by the vehicle control. ALS mice demonstrated progressive body weight loss as well, which was similarly reversed by IN-DAN treatment, but not by the vehicle control. Muscle strength, as measured by grip force, was significantly reduced in ALS mice but robustly preserved IN-DAN treatment, which prevented the decrease by 213% (P<0.0001), while the vehicle control had no effect. Spinal cord weight was significantly reduced in ALS mice, a trend reversed by intranasal dantrolene nanoparticle treatment, but not by the vehicle control. Survival analysis revealed that 100% of control mice survived through the 30-day treatment period (up to 120 days of age), while survival in untreated or vehicle-treated ALS mice dropped to 67%. In contrast, ALS mice treated with intranasal dantrolene nanoparticles demonstrated a significantly improved survival rate of 89%. Thus, intranasal dantrolene nanoparticle treatment significantly and robustly improved neurological outcomes in SOD1-G93A ALS mice, inhibiting neurological impairment, motor dysfunction, balance deficits, and muscle weakness. These improvements were associated with a marked inhibition of spinal cord weight loss. Additionally, dantrolene treatment reversed body weight loss and significantly improved survival probability in ALS mice.

## Introduction

Amyotrophic lateral sclerosis (ALS) is a fatal neurodegenerative disease characterized by the progressive degeneration of selective motor neurons, leading to muscle weakness, paralysis, and eventual death due to respiratory failure[1-3]. ALS has a prevalence of 6-9 per 100,000 people, and a lifetime risk of approximately 1 in 350 individuals. The disease carries a poor prognosis, with an average survival of 2–5 years following diagnosis. ALS is categorized into familial ALS (fALS) and sporadic ALS (sALS). fALS accounts for approximately 10% of cases and is associated with a known family history or genetic mutations, whereas sALS represents the remaining 90% of cases and typically has an unknown etiology. Pathologically, ALS is defined by the loss of upper and lower motor neuron cell bodies, as well as degeneration of the corticospinal tracts and lower motor neuron axons. These changes lead to denervation of skeletal muscle[2], resulting in muscle atrophy and progressive weakness. Unfortunately, current treatments are limited to symptom relief, and no approved disease-modifying therapies exist for ALS[2, 4, 5].

More than 150 gene mutations have been identified in ALS[5]. The most well-characterized gene mutations associated with ALS include Cu/Zn superoxide dismutase 1 (SOD1), Fused in Sarcoma (FUS), TAR DNA-binding protein 43 (TDP-43), and Chromosome 9 open reading frame 72 (C9ORF72)[6]. Approximately 20% of fALS cases involve mutations in the SOD1 gene, which encodes an enzyme responsible for converting superoxide radicals into molecular oxygen and hydrogen peroxide, scavenging harmful oxygen species that can damage cells[7]. Mutations in SOD1 impair this antioxidant function, resulting in oxidative stress, neurotoxic gain-of-function effects, and neurodegeneration[8, 9]. The SOD1-G93A gene mutation is commonly found in ALS patients, and transgenic mice carrying the SOD1-G93A mutation are the most widely used animal model of fALS for studying ALS mechanisms and pathologies, and for evaluating therapeutic efficacy[10-12]. Among many proposed pathologies in ALS, intracellular calcium (Ca^2+^) dysregulation plays a critical upstream role in motor neuron degeneration and muscle atrophy. Early upstream Ca^2+^ dysregulation triggers multiple downstream pathologies, including increased reactive oxygen species (ROS) and oxidative stress[13, 14], endoplasmic reticulum (ER) stress[15], impaired protein folding and aggregation[16], defective autophagy[14], mitochondrial damage and dysfunction[17, 18], neuroinflammation[19], disrupted nucleocytoplasmic and vesicle transport, impaired DNA damage repair, aberrant RNA metabolism, axonopathy, and motor neuron and muscle cell death through apoptosis[16, 20], pyroptosis[21, 22] and ferroptosis[23], etc. Thus, therapeutic strategies targeting and correct upstream calcium dysregulation are expected to ameliorate downstream pathologies, and therefore may offer an effective approach for treating ALS.

Physiological regulation of Ca^2+^ concentrations in the cytosol, endoplasmic reticulum (ER), and mitochondria plays a critical role in nearly all cellular functions. However, Ca^2+^ dysregulation, characterized by pathological increases in cytosolic and mitochondrial Ca^2+^ and concurrent depletion of ER Ca^2+^, contributes significantly to ALS pathogenesis[24]. Calcium dysregulation in ALS may result from pathological Ca^2+^ influx from the extracellular space into the cytosol or from excessive Ca^2+^ release from the ER. One major mechanism is Ca^2+^ influx through the pathological overactivation of N-methyl-D-aspartate (NMDAR) receptors and α-amino-3-hydroxy-5-methyl-4-isoxazoleproprionic acid receptors (AMPARs), driven by excess extracellular glutamate or glutamate excitotoxicity, a key factor in ALS-associated Ca^2+^ imbalance[24, 25]. This glutamate mediated-excitotoxicity leads to cytosolic Ca^2+^ elevation, which in turn induces excessive Ca^2+^ release from the ER via ryanodine receptor (RyR) channels[26, 27] through a mechanism known as Ca^2+^-induced Ca^2+^ release (CICR). This cascade results in mitochondrial Ca^2+^ overload[25] and oxidative stress[28], ultimately contributing to motor neuron degeneration present in ALS[29-31]. This mechanism is considered a primary driver of pathology in SOD1 mutation-related ALS[11, 32]. Up to 87% of NMDAR activation-mediated cytosolic Ca^2+^ elevation originates from Ca^2+^ release from the ER via RyRs[33], highlighting the central role of ER dysfunction in ALS pathology. Conversely, Ca^2+^ dysregulation may originate directly from ER Ca^2+^ leak. For example, SOD1 mutations increases ROS levels and oxidative stress, which in turn promote Ca^2+^ leakage from the ER through RYRs[34] or inositol 1,4,5-trisphosphate receptors (InsP_3_Rs)[35]. This results in mitochondrial Ca^2+^ overload, pathological ROS production, impaired ATP generation, and continued ER Ca^2+^ depletion, forming a vicious cycle[36] that leads to motor neuron neurodegeneration[37-39] and muscle cell death[40, 41]. Thus, antagonists of RyRs or InsP_3_Rs are expected to be neuroprotective against ALS. Recent studies support the importance of ER Ca^2+^ leak through pathologically activated RyRs in ALS, and the therapeutic potential of RyR inhibition as a disease-modifying strategy[42].

Dantrolene, a RyR inhibitor, is the only FDA-approved drug for treating malignant hyperthermia, a condition caused by overactivation of type 1 RyRs (RyR-1)[43]. Furthermore, dantrolene has been proposed as a potential therapy for various other diseases, including cardiac arrhythemias[44], cardiac hypertrophy[45], heart failure[46], asthma[47], diabetes[48, 49], and multiple neurodegenerative disorders[22, 50]. Dantrolene has demonstrated neuroprotective effects against strokes, Huntington’s disease, and Alzheimer’s disease, primarily through its inhibition of intracellular Ca^2+^ dysregulation[50-52]. Although some studies suggest that dantrolene may inhibit NMDARs as well[53, 54], these findings require further verification. Nonetheless, dantrolene has been shown to inhibit up to 87% of NMDA-mediated abnormal cytosolic calcium ([Ca^2+^]_c_) elevation in cortical neurons[33], suggesting strong therapeutic potential in ALS, due to its potent inhibition of glutamate excitotoxicity[53, 54] and inhibition of Ca^2+^ dysregulation originating from pathological Ca^2+^ release via ER-localized RYRs[55] or InsP_3_Rs[56, 57]. While dantrolene does not appear to inhibit AMPAR-mediated Ca^2+^ dysregulation[58], it may ameliorate Ca^2+^ dysregulation in ALS by inhibiting InsP_3_R[56, 57]-mediated Ca^2+^ release and blocking mitochondrial RyRs involved in ER-to-mitochondria calcium transfer[59]. One prior study demonstrated that dantrolene was neuroprotective in *in vitro* tissue models, but not in *in vivo* ALS animal models[60]. This lack of efficacy was likely due to limited central nervous system (CNS) penetration via traditional oral or intravenous routes of administration[60, 61]. In contrast, our investigations using intranasal dantrolene nanoparticles demonstrated significantly higher dantrolene concentrations and longer duration in the brain, as well as an increased brain-to-blood concentration ratio[62, 63], especially in aged mice[64], compared to oral or subcutaneous administration methods. Considering the central role of Ca^2+^ dysregulation in ALS pathology, and the enhanced CNS delivery achieved via the intranasal nanoparticle formulation, the neuroprotective potential of intranasal dantrolene nanoparticles warrants investigation. Therefore, we evaluated the therapeutic efficacy of intranasal dantrolene nanoparticles on motor dysfunction, muscle weakness, and survival probability in the most widely used ALS animal model, the SOD1-G93 transgenic mouse.

## Materials and Methods

### Animals

All procedures were approved by the Institutional Animal Care and Use Committee (IACUC) of the University of Pennsylvania. Four breeding pairs of wild-type (C57BL/6SJLF1) mice (Strain #000664) and four breeding pairs of SOD1 (SOD1*G93A) mice (Strain #004435), with the BL/6 SJLF1 background, were purchased from the Jackson Laboratory and used to establish colonies. Mice were housed in the University of Pennsylvania laboratory animal research facilities at a temperature of 21–22°C with a 12–h light-dark cycle, and were provided with food and water ad libitum and nestlets for enrichment. All mice were weaned at no later than one month of age and genetically identified via polymerase chain reaction (PCR) analysis prior to weaning. After genotyping, mice were divided into different cages according to age and gender, with no more than five mice per cage. Mice were separated if aggressive behavior or fighting was observed. Behavioral observations were conducted daily, and body weight was recorded each week. All efforts were made to minimize animal suffering and the number of mice used, in accordance with IACUC guidelines. Both male and female mice were used in this study.

### Intranasal Dantrolene Nanoparticle Preparation and Administration

Dantrolene (Sigma, St. Louis, MO) was dissolved in the Ryanodex formulation vehicle (RFV: 125 mg mannitol, 25 mg polysorbate 80, and 4 mg povidone K12, dissolved in 5 mL sterile water; final pH adjusted to 10.3), as described previously in our prior publications[62, 62]. For intranasal administration, the final concentration of dantrolene was 5 mg/mL. Mice were gently restrained by the scruff of their necks using one hand. With the other hand, 1 μL of dantrolene nanoparticle solution (or the empty vehicle control) was administered per gram of body weight. For example, a mouse weighing 20 grams received 20 μL solution, equivalent to a dantrolene dose of 5 mg/kg. The solution was administered drop by drop directly into the nostrils, allowing for gradual absorption. Special care was taken to minimize stress and ensure proper intranasal delivery to the nasal cavity, without entry into the stomach or lungs.

### Experimental Groups

As illustrated in Supplemental Figure 1, age-matched male and female mice were randomly assigned to experimental groups at the time of genotyping, around 1 month of age. Wild-type (WT) C57BL/6SJLF1 and SOD1-G93A transgenic mice were divided into three experimental groups: 1) No treatment control (NO TX), 2) Intranasal dantrolene in Ryanodex formulation vehicle (IN-DAN), and 3) Intranasal Ryanodex vehicle without dantrolene (IH-VEH; vehicle control). IN-DAN and IN-VEH treatments began at 90 days of age, administered once daily, 7 days per week, for 30 consecutive days, until the mice reached 120 days of age. Each treatment group contained 10-18 mice, for a total of 83 mice across all six experimental groups (WT control and SOD1-G93A x 3 treatment conditions). Both RFV (IN-VEH) and RFV with dantrolene (IN-DAN) were freshly prepared before each treatment, following our previously published protocols[62, 63]. All behavioral testing was performed immediately after completion of the 30-day treatment period, except for the rotarod test, which was conducted weekly during the 30-day treatment window. Mice were euthanized following the conclusion of behavioral assessments.

### Behavioral Tests

#### Rotarod Test

To evaluate the progressive changes in motor coordination and balance, the rotarod test was performed once weekly during the treatment period, using a method previously described[63]. The latency to fall from an accelerating rotarod (IITC Series 8, Life Sciences, Woodland Hills, CA) was measured as an indicator of motor function. In brief, mice were acclimated to the testing room for at least one hour prior to testing. Animals underwent two 60-second training trials, separated by a 30-minute rest period. Following training, three 120-second test trials were conducted. Each test trial involved gradual acceleration from 4 to 40 rpm, and trials were spaced 60 minutes apart to ensure adequate recovery and limit stress. Latency to fall was recorded and used as a measure of motor performance, with shorter durations indicating impaired motor coordination and balance.

#### Balance Beam Test

Following the completion of the 30-day treatment period, the balance beam test was performed to assess motor coordination and balance. As shown in Supplemental Video 1 (https://drive.google.com/file/d/14iJB82VOsxZOjKOhDzqY532YrL96PPXM/view?usp=sharing), an 80 cm-long rectangular wooden beam, with a width of either 12 or 6 mm, was used. Each beam was securely mounted between two vertical poles, positioned 50 cm above the ground. Padding was positioned underneath the beam to protect mice from injury in the event of falls and to collect droppings. A black box containing food was placed at one end of the beam to serve as the target area and incentive for traversal. Mice were placed at the starting end of the beam, and the time required to traverse the full 80 cm was recorded. A maximum time limit of 3 minutes was imposed. If a mouse stalled on the beam, a small amount of food was placed midway to encourage continued movement. After each trial, the beam was cleaned of droppings and wiped down with 70% ethanol, followed by water. Each mouse underwent three training trials on each of the two beam widths, for a total of six training trials, with 30-minute rest intervals provided between sessions. Following training, one testing trial was conducted on each beam. During testing, both the time taken to traverse the beam and the number of foot slips off the beam were recorded and analyzed. Longer traversal times and a higher number of foot slips indicated poorer motor coordination and balance.

### Kondziela’s Inverted Screen Test

Following the completion of the 30-day treatment period, the inverted screen test was performed to evaluate muscle strength. As shown in Supplemental Video 2 website (https://drive.google.com/file/d/1frY2tBQCcZUODTSGzwTnT_edRs_kTwCb/view?usp=sharing), we used a 43 cm x 43 cm wire mesh screen, consisting of 12 mm square openings of 1 mm diameter wire. During testing, the screen was held over a cardboard box lined with padding, which served to collect droppings and prevent injury in the event of falls. Each mouse was placed in the center of the screen, which was then lifted to a height of 40-50 cm above the box. The screen was gradually rotated to an inverted position over 2 seconds, with the head-facing side of the mouse tilted downward first. After the screen was fully inverted, the latency to fall was recorded, with a maximum cutoff time of 2 minutes. Once either the time limit was reached or the mouse fell, the duration of time held on the screen was recorded. Three trials were conducted per mouse, with a 30-minute resting interval provided in between trials. The average latency to fall across the three trials was calculated and used for analysis. Shorter durations indicate reduced grip strength and impaired muscle function.

### Neurological Scoring

The neurological score (NS) was used to evaluate overall neurological function, as an indicator of ALS disease progression. Scores were assessed independently for each hind limb of each mouse, based on the combined results of three functional tests, following a modified protocol described in Supplemental Table 1[65]: 1) Tail suspension test, 2) Walking gait observation, and 3) Righting reflex test. To perform the tail suspension test, each mouse was gently lifted approximately 3 cm from the base of its tail and suspended above its home cage for 2 seconds. This was repeated twice more, for a total of three trials. Hind limb movement was observed, with the most consistent outcome recorded. For the walking gait assessment, the mouse was placed on a clean flat surface and allowed to walk a distance of 75 cm. Gait and posture were observed and documented (refer to Supplemental Video 3 website for open field activity: https://drive.google.com/file/d/100QVvRzTwBDF5XFyfunhtSXp7LVaEbC6/view?usp=sharing). To conduct the righting reflex test, the mouse was placed recumbent on either its left or right side. The time required for the mouse to right itself unassisted was measured. Upon completion of all three tests, neurological scores for each hind limb were assigned independently on a scale from 0 to 4, based on the criteria outlined in Supplemental Table 1. A neurological score of 0 (NS = 0) reflects a normal neurological function, while a score of 4 indicates severe impairment or end-stage disease, with scores reflecting the overall motor performance and neurological status of each mouse.

### Euthanasia and Tissue Collection

Mice from all experimental groups were euthanized at approximately 135 days of age, following completion of all behavioral tests (see Supplemental Figure 1). As previously described[63, 64], animals were deeply anesthetized with 2-4% isoflurane delivered via nose cone, with the concentration adjusted based on responsiveness to a toe pinch. After confirming deep anesthesia, the skin was prepped and an incision was made to open the chest and expose the heart. Blood was collected directly from the heart using a 27G needle and heparinized syringe for serum and plasma analysis. Blood samples were centrifuged at 1,400 RPM at 4°C for 30 minutes, and the plasma supernatant was collected and frozen at -80°C. Mice were euthanized via transcardial perfusion with ice-cold phosphate-buffered saline (PBS) at 4 ºC, followed by exsanguination. The brain and spinal cord were carefully harvested. The right hemisphere of the brain was dissected, and relevant regions (e.g., cortex) were fresh-frozen at -80° C for subsequent protein analysis via western blotting. The left hemisphere was fixed in 4% paraformaldehyde, followed by paraffin embedding for histological or immunohistochemical studies. The spinal cord was bisected at approximately the T12 or L1 level. The upper half (C1-T12) was fixed in 4% paraformaldehyde for histology and immunohistochemistry, while the lower half (L1-S5) was fresh-frozen at -80ºC for immunoblotting studies.

### Protein Level Determination by Immunoblotting (Western Blot)

Protein expression levels were measured using immunoblotting (western blot analysis). Tissue samples from the lower half of the spinal cord were extracted and homogenized in ice-cold RIPA buffer (9806S, Cell Technology, USA), supplemented with protease inhibitor cocktails (P8340, Roche). Homogenates were rocked at 4 °C for 90 min, then centrifuged at 14,000 rpm using a Denville 260D brushless microcentrifuge for 20 minutes at 4 °C to remove cellular debris. The supernatant was collected, and protein concentrations were determined using the BCA Protein Assay Kit (Pierce, Rockford, IL, USA). Equal amounts of protein (50 µg/lane) were loaded onto 4–20% TGX precast mini-protein gels (Cat. #4,561,094, BIO-RAD), followed by electrophoretic transfer to polyvinylidene difluoride (PVDF) membranes (Immobilon-P, MERK Millipore, Ireland) using a wet electrotransfer system (BIO-RAD, USA). After transfer, membranes were blocked in 5% BSA (Sigma-Aldrich) for 1 hour, then incubated overnight at 4 °C with primary antibodies, including those targeting neurofilament light chain (NFL). The following day, membranes were incubated with HRP-conjugated secondary antibodies, including anti-mouse IgG1 and anti-rabbit IgG, then washed with tris-buffered saline containing 0.2% Tween-20 (TBST). Protein bands were visualized using ECL Western Blotting Detection Reagents (Cytiva, Amersham, UK), and band intensity was quantified using ImageJ software (National Institutes of Health, Bethesda, MD, USA).

### Spinal Cord Weight Measurement

The weights of the upper and lower halves of the spinal cord were measured using an AG104 balance (Mettler-Toledo, Mumbai, India). Care was taken to preserve the integrity of the spinal cord during its dissection and isolation from the vertebral column.

### Statistics

All data are presented as mean ± standard error of the mean (SEM). Statistical analyses were performed using GraphPad Prism software (version 9.3.1, GraphPad Software, CA, USA). Comparisons involving two genotypes of mice, and two treatment groups were analyzed using two-way analysis of variance (ANOVA), followed by Tukey’s multiple comparisons test. A p value < 0.05 was considered statistically significant.

## Results

### Intranasal dantrolene nanoparticles robustly and significantly inhibited overall neurological dysfunction in SOD1-G93A transgenic ALS mice

The neurological scoring protocol is commonly used to monitor disease progression and neurological function in SOD1-G93A mice[65]. We assessed neurological scores at 120 days of age, following completion of the 30-day treatment period. Higher scores reflect worse neurological function and greater ALS severity. In SOD1-G93A transgenic ALS mice, intranasal dantrolene nanoparticle (IN-DAN) robustly and significantly reduced neurological score by 90%, in comparison to mice that underwent no treatment (3.167 vs. 0.313, P<0.0001). Although intranasal vehicle (IH-VEH) treatment demonstrated some reduction in neurological score, its effect was substantially less potent than that of IN-DAN. Importantly, in non-transgenic control mice, neither IN-DAN nor IH-VEH produced significant changes in neurological score, indicating the treatment possessed no off-target neurological effects in healthy animals. Supplemental Video 3 website demonstrates overall observed behavioral activity in mice from different experimental groups within an open field environment (https://drive.google.com/file/d/100QVvRzTwBDF5XFy-funhtSXp7LVaEbC6/view?usp=sharing).

### Intranasal dantrolene nanoparticles robustly and significantly inhibited impairment of motor coordination and balance in SOD1-G93A transgenic ALS mice

We used beam balance tests with either a 12 mm-wide beam or a 6 mm-wide beam to assess motor coordination and balance, following a previously described method[66], as demonstrated in the supplemental video 2 website (https://drive.google.com/file/d/14iJB82VOsxZOjKOhDzqY532YrL96PPXM/view?usp=sharing). Increased time to cross the beam and a greater number of foot slips indicate more severe motor impairment. Compared to non-transgenic control mice, SOD1-G93A mice required significantly more time to cross both the 12 mm and 6 mm beams, with 10.78-fold (116.9 vs. 9.92 seconds, P<0.0001) and 5.27-fold (112.2 vs. 17.90 seconds, P<0.0001) increases, respectively (Fig. 2 A, C). IN-DAN treatment significantly reduced crossing times for the SOD1-G93A mice on both beams, by 78% (116.9 vs. 25.66 seconds, P<0.0001) on the 12 mm beam and 79% (112.2 vs 33.19 seconds, P<0.0001) on the 6 mm beam. While intranasal vehicle (IH-VEH) treatment also significantly reduced crossing time on the 6 mm beam (112.2 vs. 50.90 seconds, P=0.0034), it did not significantly affect performance on the 12 mm beam. Similarly, compared to controls, SOD1-G93A mice exhibited a dramatic increase in the number of foot slips while crossing the 12 mm and 6 mm beams, 33.7-fold (0.250 vs. 8.67, P<0.0001) and 10.75-fold (0.83 vs. 9.750, P<0.0001), respectively. This impairment was significantly and robustly reduced by IN-DAN treatment, which decreased foot slips by 84% (8.67 vs. 1.38, P<0.0001) on the 12-mm balance beam and 73% (9.75 vs. 2.63, P<0.0001) on the 6-mm balance beam (Fig. 2 B, D). Although IN-VEH treatment resulted in a reduction in foot slips on both beams, the effect was not statistically significant (Fig. 2B, 2D). Supplemental Video 1 website shows representative trials on the 12 mm beam balance test across experimental groups.

**Figure 1.**
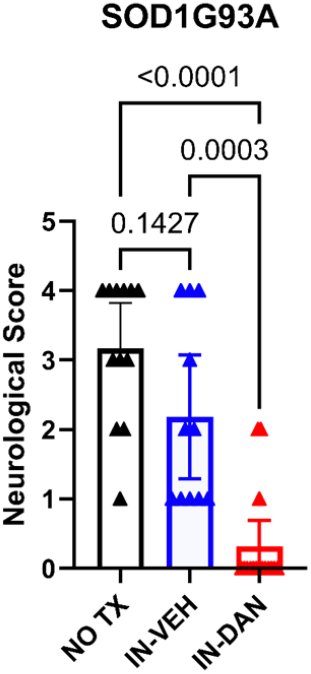
Intranasal dantrolene nanoparticles robustly and significantly improve overall neurological function in SOD1-G93A mice. Overall neurological scores were evaluated at the conclusion of a 30-day treatment period with intranasal dantrolene nanoparticles (IN-DAN), vehicle control (IN-VEH), and no treatment (NO TX) in SOD1-G93A transgenic ALS mice at 120 days of age. Higher neurological scores indicate more severe neurological function impairment. N=12 (NO TX), 10 (IN-VEH), and 16 (IN-DAN). Data are presented as means ± 95% CI and were analyzed using one-way ANOVA, followed by Tukey’s multiple comparison test (MCT). P<0.05 was considered statistically significant.

**Figure 2.**
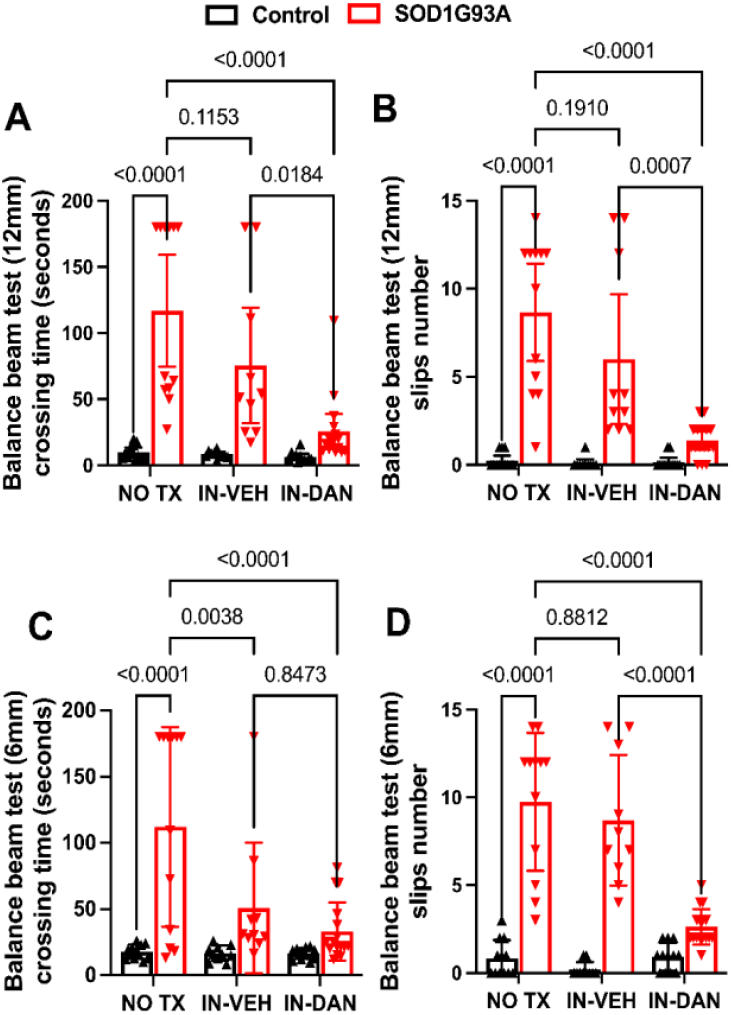
Intranasal dantrolene nanoparticles robustly and significantly inhibit impairments in motor coordination and movement balance in SOD1-G93A ALS mice. Overall motor coordination and balance were evaluated at the end of a 30-day treatment period (No treatment (NO TX), intranasal vehicle (IN-VEH) or intranasal dantrolene nanoparticles (IN-DAN) at 120 days of age, using the beam balance test at two beam widths: 12 mm (**A, B**) and 6 mm (**C, D**). Longer beam crossing time (**A, C**) and increased number of foot slips (**B, D**) indicate greater impairment of motor coordination and movement balance. N = 12 (NO TX), 10 (IN-VEH), and 12 (IN-DAN) for the non-transgenic control mice and 12 (NO TX), 10 (IN-VEH), and 16 (IN-DAN) for the transgenic SOD1-G93A ALS mice. Data are presented as means ± 95% CI and were analyzed using two-way ANOVA, followed by Tukey’s multiple comparison test (MCT). P<0.05 was statistically significant.

Furthermore, we employed a second method, the rotarod test, to evaluate motor coordination and balance in mice. This test was conducted weekly over the 30-day period of treatment (see Supplemental Fig. 2). Longer latency to fall from the accelerating rod indicates better motor coordination and balance. Compared to control wild-type (WT) mice, SOD1-G93A transgenic mice exhibited significantly reduced latency to fall across all four weeks of testing (Supplemental Fig. 2A-D). The degree of statistical significance increased progressively from week one to week four, indicating a gradual decline in motor function over time in the ALS transgenic mice. While intranasal dantrolene nanoparticle treatment (IN-DAN) showed a trend towards increased latency on the rotarod, this effect did not reach statistical significance at any time point during the 30-day treatment period.

### Intranasal dantrolene nanoparticles significantly inhibited body weight loss and muscle weakness in SOD1-G93A transgenic ALS mice

Body weight was monitored weekly throughout the treatment period. As shown in Figure 3A, non-transgenic control mice displayed a consistent increase in body weight over the four-week treatment period. In contrast, SOD1-G93A transgenic ALS mice, whether untreated or treated with intranasal vehicle (IN-VEH), exhibited significant body weight loss. Intranasal dantrolene nanoparticle treatment (IN-DAN) completely reversed this weight loss in SOD1-G93A mice, however, resulting in a body weight trajectory comparable to that of non-transgenic controls. Weight loss in SOD1-G93A transgenic mice is likely attributable to skeletal muscle atrophy, a common symptom in human ALS patients (see Supplemental Video 3 Activity Observation (https://drive.google.com/file/d/100QVvRzTwBDF5XFyfunhtSXp7LVaEbC6/view?usp=sharing). To assess muscle strength, we employed Kondziela’s inverted screen test, which measures the duration a mouse can hang from an inverted metal screen. Longer latency indicates greater grip force, or stronger muscle strength (Fig. 3 B). Compared to non-transgenic control mice, SOD1-G93A mice showed a dramatic 4.52-fold reduction in hang time (108.2 vs. 19.61 seconds, P<0.0001), indicating development of significant muscle weakness. IN-DAN treatment significantly improved grip strength, increasing hang time by 2.13-fold (19.61 vs. 61.37, P=0.0026). While IH-VEH treatment showed a trend toward improvement, the change was not statistically significant (P = 0.18). Supplemental Video 2 website displays representative trials from each experimental group (https://drive.google.com/file/d/1frY2tBQCcZUODTSGzwTnT_edRs_kTwCb/view?usp=sharing).

**Figure 3.**
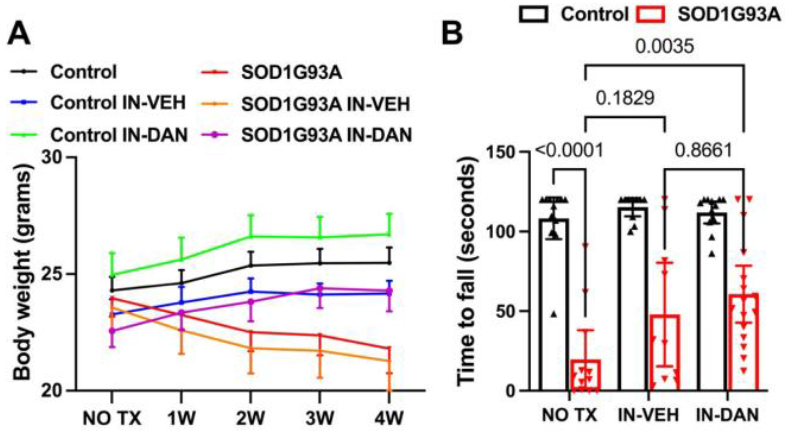
Intranasal dantrolene nanoparticles significantly inhibit body weight loss and muscle weakness in SOD1-G93A transgenic ALS mice. **A**. Body weight changes were monitored weekly (W) throughout the 4-week treatment (TX) period. **B**. Muscle strength, assessed via grip force, was evaluated at the end of the 30-day treatment period, including the no treatment (NO TX), intranasal vehicle (IN-VEH) or intranasal dantrolene nanoparticles (IN-DAN) condition groups, using the Kondziela inverted screen test. A longer latency to fall from the inverted screen indicates greater muscle strength. N=12 (NO TX), 11 (IN-VEH), 12 (IN-DAN) for non-transgenic control mice. N=12 (No TX), 10 (IN-VEH), and 16 (IN-DAN) for the transgenic SOD1G93A transgenic ALS mice (**B**). Data are presented as means ± 95% Cl and were analyzed using two-way ANOVA, followed by Tukey’s multiple comparison test (MCT). P<0.05 was considered statistically significant (**B**).

### Intranasal dantrolene nanoparticles significantly inhibited spinal cord weight loss in SOD1-G93A transgenic ALS mice

We assessed NFL levels in the lumbar and sacral spinal cord (approx. L1-S5). In SOD1-G93A mice, NFL levels were significantly decreased by 3.9 fold (1.47 vs. 0.30, P<0.0001) compared to control mice (Fig. 4 A, B), implying neurodegeneration consistent with other studies^3^. IN-DAN treatment tended to increase spinal NFL protein levels, but not to a statistically significant degree (Fig. 4C). The weight of 4% paraformaldehyde-fixed cervical and thoracic spinal cord (approx. C1-T12) was significantly reduced by 43% in SOD1-G93A mice (73.10 vs. 41.62 mg, P=0.0008), compared to the controls. This loss was significantly ameliorated by IN-DAN treatment, with spinal cord weight increasing by 53% (41.62 vs 63.60 mg, P<0.019), but not by the IN-VEH treatment (Fig. 4D, 4E). Similarly, the weight of fresh lumbar and sacral spinal cord (approx. L1-S5) was reduced by 71% (30.99 vs 8.90 mg, P<0.0001) in SOD1-G93A mice compared with non-transgenic controls. IN-DAN treatment significantly restored spinal cord weight, increasing it by 1.43-fold (8.90 vs. 21.66 mg, P=0.034), while IH-VEH had no significant effect (Fig. 4F).

**Figure 4.**
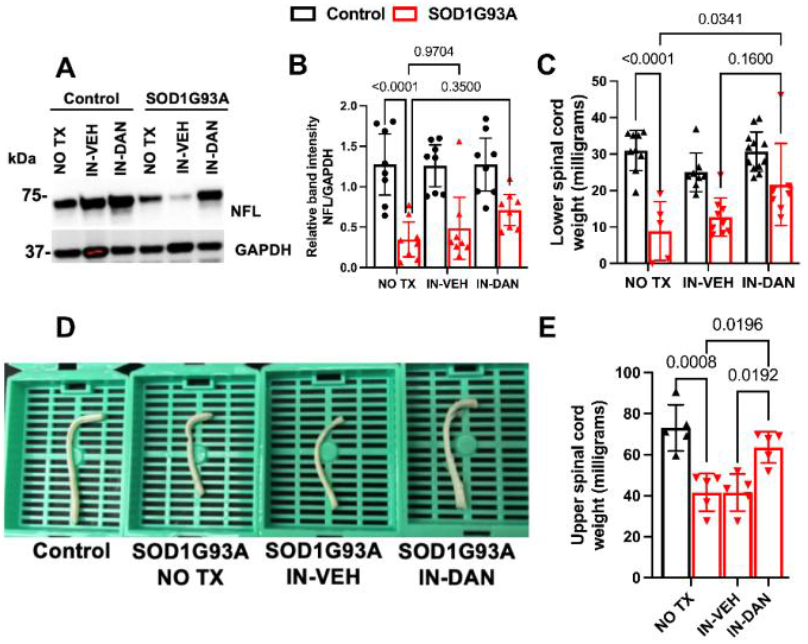
Intranasal dantrolene nanoparticles significantly inhibit spinal cord degeneration in SOD1-G93A ALS mice. Spinal cord degeneration was evaluated following 30 days of treatment with no treatment (NO TX), intranasal vehicle control (IN-VEH) or intranasal dantrolene nanoparticles (IN-DAN), at 135 days of age, by measuring levels of the neuronal skeletal protein neurofilament light chain (NFL) and spinal cord weight. (**A**) Representative Immunoblot illustrating changes in NFL protein level in the lumbar and sacral spinal cord (∼L1 to S5). (**B)** Quantification and statistical analysis of NFL protein expression in the same region. N = 8 per group in both control and SOD1-G93A mice. (**C**) Fresh weight of lumbar and sacral spinal cord tissue collected for Western blot (∼L1 to S5). N=10 per group in control mice, N=5 (NO TX), 9 (IN-VEH) and 7 (IN-DAN) in SOD1-G93A transgenic mice. **D**. Representative images of 4% paraformaldehyde-fixed cervical and thoracic spinal cords (∼C1 to T12). **E**. Weight measurements of the cervical and spinal cord segments (∼C1 to T12). N = 5 per experimental group. Data are presented as means ± 95% Cl and were analyzed using two-way ANOVA, followed by Tukey’s multiple comparison test (MCT). (**B, C, E**). P<0.05 was considered statistically significant.

### Intranasal dantrolene nanoparticles robustly prolonged survival probability in SOD1-G93A transgenic ALS mice

Neither intranasal dantrolene nanoparticle (IN-DAN) nor vehicle (IH-VEH) treatments affected survival in non-transgenic control mice, which maintained 100% survival throughout the 30-day treatment period (data not shown). In contrast, in SOD1-G93A transgenic ALS mice, survival declined from 100% at the treatment onset to 67% by the end of the 30-day period, in both untreated and IH-VEH treated groups. Strikingly, however, IN-DAN treatment significantly improved survival, with 89% of SOD1-G93A mice surviving through the full treatment course (Fig. 5), suggesting a robust neuroprotective effect of intranasal dantrolene nanoparticles.

**Figure 5.**
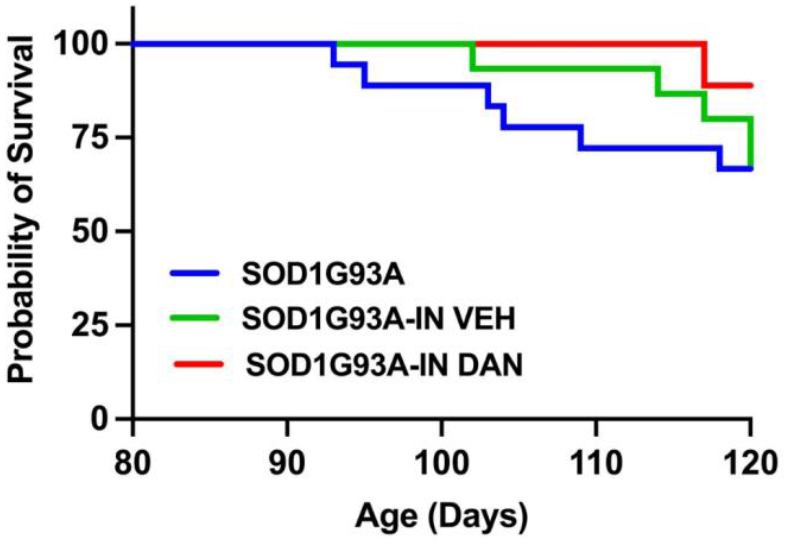
Intranasal dantrolene nanoparticle treatment robustly prolongs survival probability in SOD1-G93A transgenic ALS mice. The probability of survival in SOD1-G93A transgenic ALS mice was monitored daily over a 30-day treatment period with intranasal dantrolene nanoparticle (IN-DAN), vehicle control (IN-VEH), or no treatment (NO TX), beginning at 90 days of age and ending at 120 days. At treatment onset, N = 18 (NO TX), 15 (IN-VEH), and 18 (IN-DAN) for SOD1-G93A mice. Kaplan-Meier survival analysis was performed to evaluate survival probability. At the end of the treatment period, the survival rate for all non-transgenic control mice was 100% (data not shown). In contrast, among SOD1-G93A mice: No TX: 67% survival (12/18), IN-VEH: 67% survival (10/15), IN-DAN: 89% survival (16/18).

## Discussion

This study has demonstrated that intranasal dantrolene nanoparticles provide robust and significant therapeutic effects against motor neuron dysfunction, motor discoordination, balance impairments, muscle weakness, and reductions in body and spinal cord weight, while prolonging survival in an ALS mouse model carrying the SOD1-G93A mutation. These beneficial effects were observed even when treatment was initiated after the full onset of disease pathology and symptoms, at 3 months of age, suggesting dantrolene might function as a potential disease-modifying therapy of ALS.

Two primary symptom-relief drugs currently used to treat ALS patients are riluzole and edaravone. Riluzule acts as an NMDAR antagonist and edaravone functions as a scavenger of reactive oxygen species[68]. However, there remains an urgent need to develop effective disease-modifying therapies. Repurposing clinically available, FDA-approved drugs with known safety profiles and minimal organ toxicity, such as dantrolene, offers a promising strategy to accelerate clinical translation. This new formulation of dantrolene, Ryanodex, consists of crystalline nanoparticles[70]. While Ryanodex is FDA-approved for the treatment of malignant hyperthermia, its current application is intravenous, not intranasal[70]. A prior study failed to demonstrate efficacy of dantrolene in the same SOD1-G93A model[60], most likely due to its limited ability to cross the blood-brain barrier (BBB)[71]. Intranasal delivery of drugs, particularly in nanoparticle forms, has been well established to enhance BBB penetration, increasing drug concentration in the CNS and its retention time, thereby improving therapeutic efficacy while minimizing peripheral side effects and organ toxicity. This strategy is especially advantageous for treatment of long-term conditions, such as those involving chronic neurological diseases[72]. In fact, intranasal administration of riluzole nanoemulsions has been shown to produce higher brain concentrations than standard oral administration, enhancing its therapeutic potential in ALS[73]. Consistently, our previous studies demonstrated that intranasal administration of dantrolene nanoparticles in the Ryanodex formulation significantly increased brain concentration, prolonged drug duration, and enhanced the brain-to-blood concentration ratio compared to oral or subcutaneous administrations[62-64]. Intranasal dantrolene nanoparticles administered at the same dose (5mg/kg), daily, 5 days per week for up to 12 months, significantly inhibited memory loss in the 5XFAD Alzheimer’s disease mouse model, without evidence of side effects or nasal/liver toxicity[63, 74]. Given the chronic nature of ALS, the low systemic toxicity and high CNS penetration of intranasal dantrolene nanoparticle treatment further favor its utility for long-term treatment. In comparison with prior studies, the robust and significant neuroprotective and muscle-preserving effects observed in SOD1-G93A mice suggest enhanced drug delivery to the brain and spinal cord, with promoted efficacy in the CNS. Moreover, because ALS patients frequently develop difficult swallowing, intranasal drug delivery may be a more convenient and patient-friendly route of administration. A significant proportion of ALS patients develop cognitive impairment and depression-related psychiatric symptoms[75]. Accordingly, as intranasal dantrolene nanoparticles have demonstrated efficacy in treating memory dysfunction[63] and depression- and anxiety-like behaviors[76], this approach may offer additional clinical benefit beyond motor function preservation in the treatment of ALS. Similar to our previous findings in 5XFAD mice[63], the intranasal administration of vehicle alone (only the Ryanodex formulation) demonstrated predominantly minor, non-significant trends towards improvement in motor function, balance, and muscle strength, but to a clearly less effective degree than dantrolene-loaded nanoparticles. This suggests that dantrolene is the primary active agent providing neuroprotection and preserving muscle strength, although the potential additive role of the vehicle alone warrants further investigation.

Besides the general side effeccts of dantrolene such as nausea and vomitting and spleepness[80], some specific side effects or organ toxicity with chronid use of dantrolene may be concered when it is proposed to treat ALS patients. For example, chronic administration of dantrolene has been demostrated to cause plual effusion in some rear cases[81, 82]. A sever ortan toxiciy after chronic use of dantrolene at high dose for long duration can cause liver toxicity, which typically happen in female adult older than 35 years old, so the liver function need to be examined periodically if dantrolene is used to treat ALS patients in the future[80]. Intranasal dantrolenen nanoparticles for up to 10 months dis not cuase liver structure and function changes in 5XFAD mice[63, 74]. Although there has been consern to use dantrolene in ALS patients with muscle weakness as dantrolene is considered a muscle relaxant, it is an FDA approved drug to treat muscle spasm due to upper motor dysfunction, spincal cord injury, multiple sclerosis, cerebral palsy and stroke etc [83]. We speculate that the early use of dantrolene increase muscle strength by inhibiting the root cause (degeneration of motor neuron) as demonstrated in this study, while the late use of dantrolene for ALS patients with existing respiratory failure may need particular caution for potential deteriation of respiratory failure by dantrolene due to its effects to relaxm respiratory muscle, which need future clinical studies. Although dantrolene is typically considered a muscle relaxant, intranasal dantrolene nanoparticles for up to 10 months treatment did not impair muscle strenth in 5XFAD mice[63]. In this ALS model, it actually significantly reversed muscle weakness and improve muscle strengh, likely due to its inhition of motor neuron degeneration[60, 84], and potentially the direct protection against muscle damage[85]. Overall, this preclinical study strongly supports the need for urgent clinical trials to evaluate the therapeutic potential of intranasal dantrolene nanoparticles in ALS patients.

The robust inhibition of reduction in spinal cord weight observed in this study suggest that intranasal dantrolene nanoparticles inhibit degeneration of spinal cord, although the specific neuronal populations affected remain to be clarified. Although our data strongly suggest protection in the spinal cord, the exact neuroprotective mechanisms warrant further investigation. Dantrolene has previously been shown to inhibit neuronal death through multiple pathways, including apoptosis[77], pyroptosis[78], and ferroptosis[79]. At the molecular level, its ability to inhibit upstream intracellular Ca^2+^ dysregulation, triggered by pathological activation of NMDAR[33], RyR[55] and InsP_3_R[56, 57], may contribute significantly to its broad neuroprotective effects. However, further studies are needed to investigate these mechanisms in ALS pathology.

Intranasal dantrolene nanoparticles prolonged survival in SOD-G93A mice, even with a limited 30-day treatment period (from postnatal day 90 to 120). Since most ALS patients succumb to respiratory failure due to respiratory muscle weakness, the observed inhibition of spinal cord degeneration and muscle atrophy by dantrolene likely contributed to the increased survival probability. We speculate that continued treatment beyond 120 days would further extend survival in this ALS animal model.

Although this pioneering study provides compelling evidence, several limitations should be recognized and guide future investigations: 1). Relatively low sample sizes (approximately n=10 per group) in some behavioral experiments; 2) Dose-response and time-course studies are needed to determine the minimum effective dose and assess the drug’s ability to prevent, halt, or delay disease progression; 3). The precise molecular mechanisms by which dantrolene confers neuron and muscle protection in ALS remain to be elucidated, especially in the examination of degeneration of motor neurons in brain and spinal cord; 4). Potential side effects and systemic toxicity associated with long-term treatment were not evaluated in this study.

## Conclusion

In conclusion, intranasal dantrolene nanoparticles provided robust and significant improvements in overall disease severity, including motor discoordination, imbalance, muscle weakness, body and spinal cord weight loss, and survival in SOD-G93A ALS mice. These findings support further preclinical and clinical studies to evaluate intranasal dantrolene nanoparticles as a future effective disease-modifying treatment for ALS patients.

## Acknowledgements

The research was conducted in the laboratory of Dr. Huafeng Wei and should be attributed to the Department of Anesthesiology, University of Pennsylvania.

## Author Contributions

H.W. conceived and designed the study. P.B., Y.Y., B.W., A.Y., L.D., and H.W. conducted experiments and acquired the data. P.B., A.Y., and H.W. analyzed data. H.W. and Y.Y. wrote the manuscript. All authors reviewed and approved the final manuscript.

## Data Availability

Data is available upon request.

---

**Supplemental Table 1.**
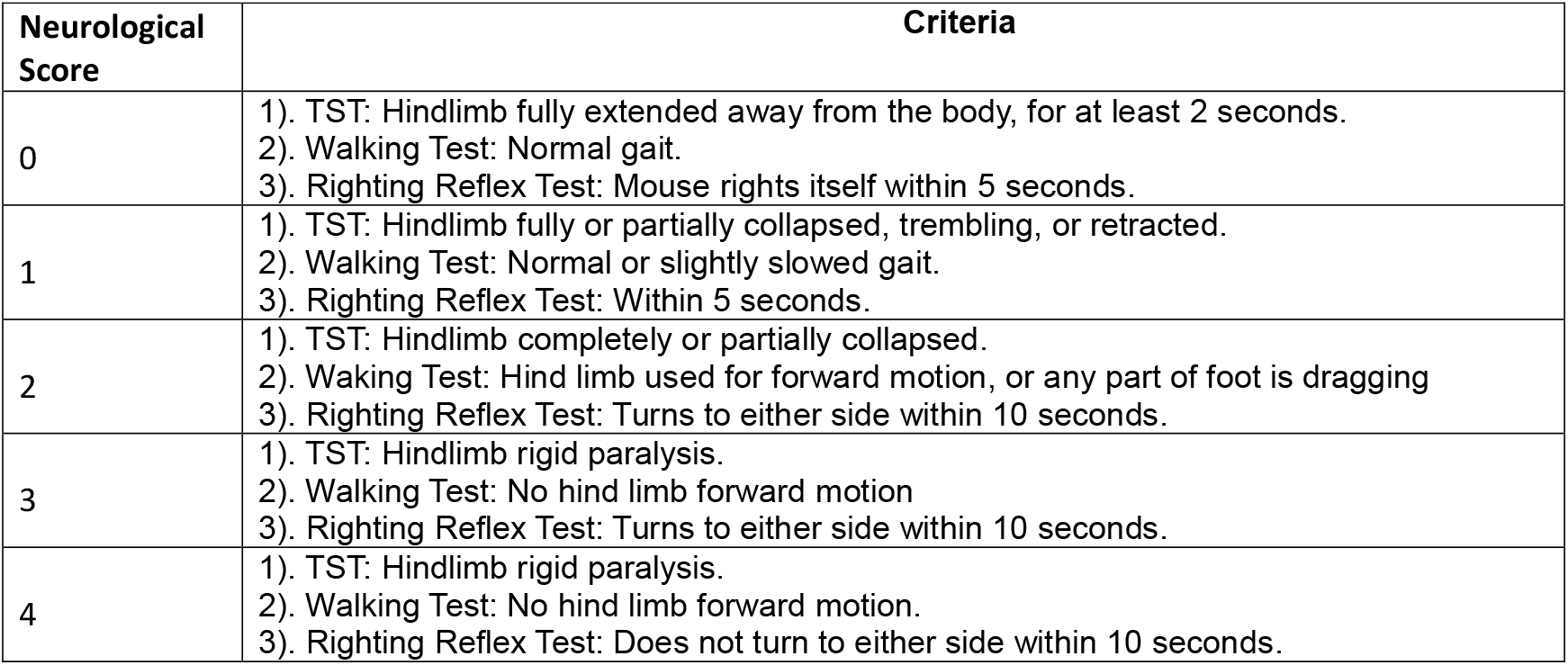
Criteria for Neurological Scores.

**Supplemental Figure 1.**
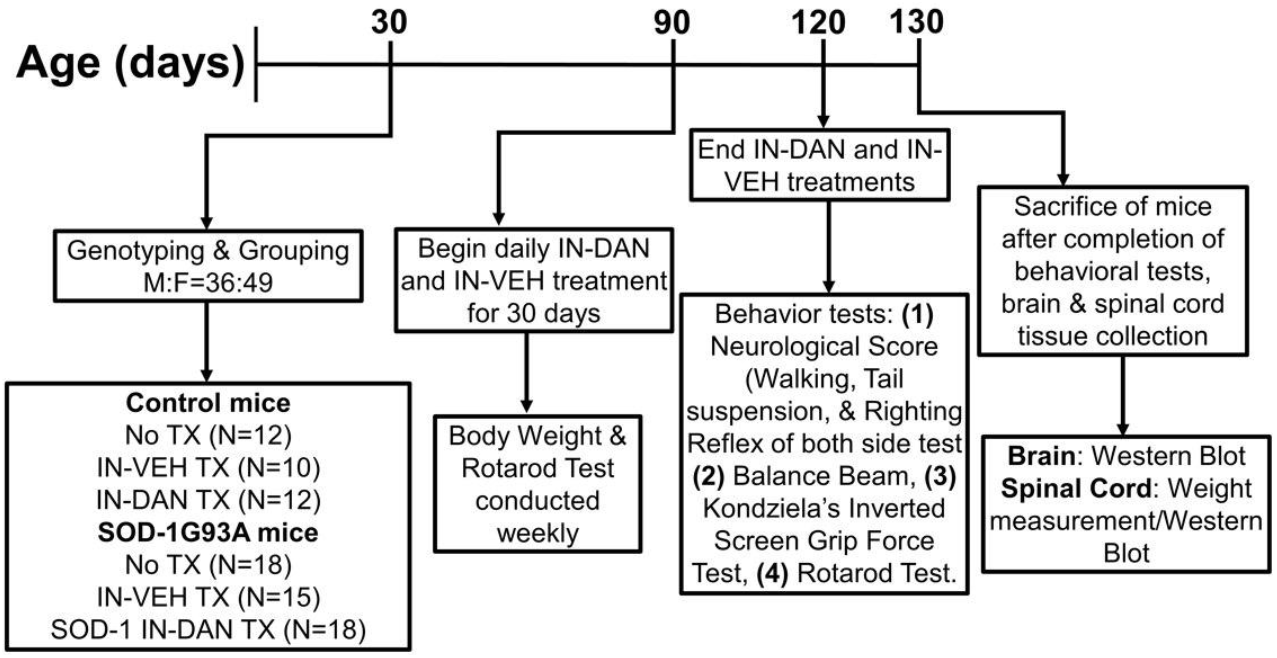
Experimental Design. No TX: No treatment. IN-DAN: Intranasal dantrolene nanoparticle treatment. IN-VEH: Intranasal vehicle control.

**Supplemental Figure 2.**
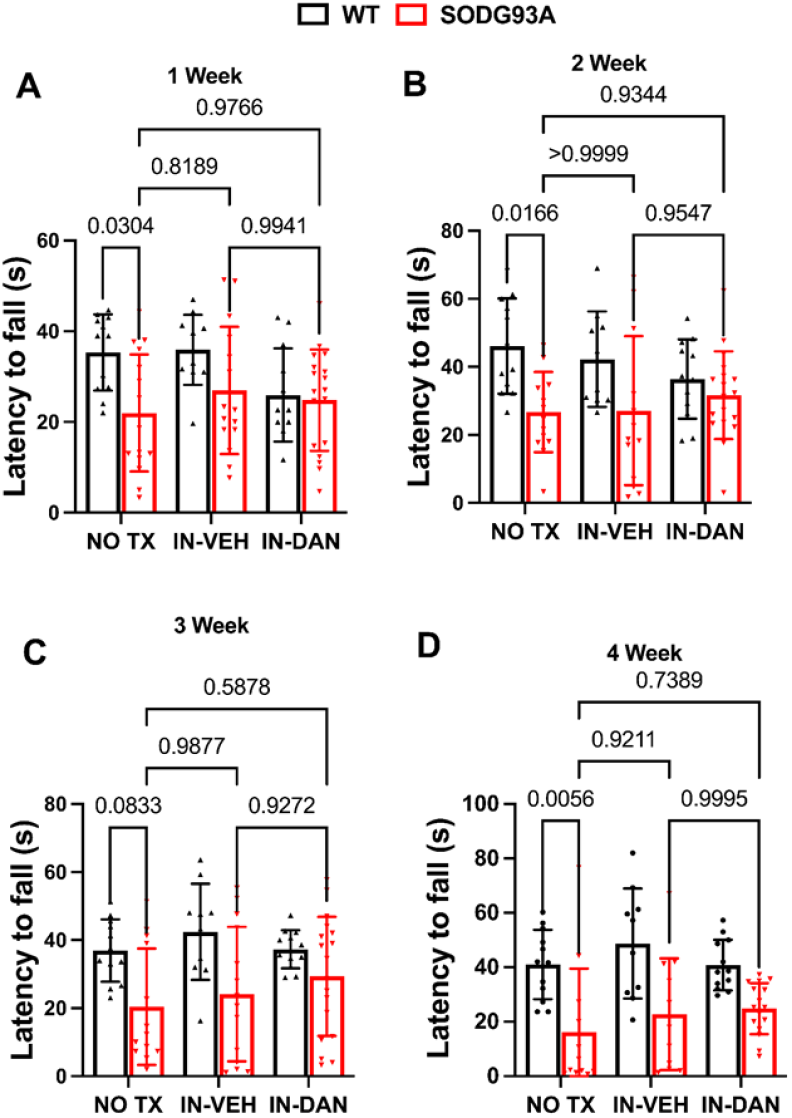
Effects of intranasal dantrolene nanoparticles on motor coordination and balance. Motor coordination and balance were evaluated weekly over a four-week treatment period using the rotarod test, following the initiation of intranasal dantrolene particle treatment (IN-DAN), vehicle control (IN-VEH), or no treatment (NO TX) between the ages of 90 and 118 days. A longer latency to fall from the rotarod indicates improved motor coordination and balance. At treatment initiation, N = 12 (NO TX), 10 (IN-VEH) and 12 (IN-DAN) for the non-transgenic control mice. N = 18 (No TX), 15 (IN-VEH) and 18 (IN-DAN) for the SOD1-G93A transgenic ALS mice. Data are presented as means ± 95%Cl and were analyzed using two-way ANOVA, followed by Tukey’s multiple comparison test (MCT).

**Supplemental Video 1. Beam Balance Test. Website:**

https://drive.google.com/file/d/14iJB82VOsxZOjKOhDzqY532YrL96PPXM/view?usp=sharing

**Supplemental Video 2. Grip Strength Test on Inverted Screen. Website:**

https://drive.google.com/file/d/1frY2tBQCcZUODTSGzwTnT_edRs_kTwCb/view?usp=sharing

**Supplemental Video 3. Activity Observation. Website:**

https://drive.google.com/file/d/100QVvRzTwBDF5XFyfunhtSXp7LVaEbC6/view?usp=sharing

