## Supplemental Data and Video Files for "Intranasal Dantrolene Nanoparticles for Treatment of Amyotrophic Lateral Sclerosis as a Disease-Modifying Drug"

### Supplemental Data File

**Supplemental Table 1. Criteria for Neurological Scores**

| Neurological Score | Criteria |
|---|---|
| 0 | 1). TST: Hindlimb fully extended away from the body, for at least 2 seconds.<br>2). Walking Test: Normal gait.<br>3). Righting Reflex Test: Mouse rights itself within 5 seconds. |
| 1 | 1). TST: Hindlimb fully or partially collapsed, trembling, or retracted.<br>2). Walking Test: Normal or slightly slowed gait.<br>3). Righting Reflex Test: Within 5 seconds. |
| 2 | 1). TST: Hindlimb completely or partially collapsed.<br>2). Waking Test: Hind limb used for forward motion, or any part of foot is dragging<br>3). Righting Reflex Test: Turns to either side within 10 seconds. |
| 3 | 1). TST: Hindlimb rigid paralysis.<br>2). Walking Test: No hind limb forward motion<br>3). Righting Reflex Test: Turns to either side within 10 seconds. |
| 4 | 1). TST: Hindlimb rigid paralysis.<br>2). Walking Test: No hind limb forward motion.<br>3). Righting Reflex Test: Does not turn to either side within 10 seconds. |

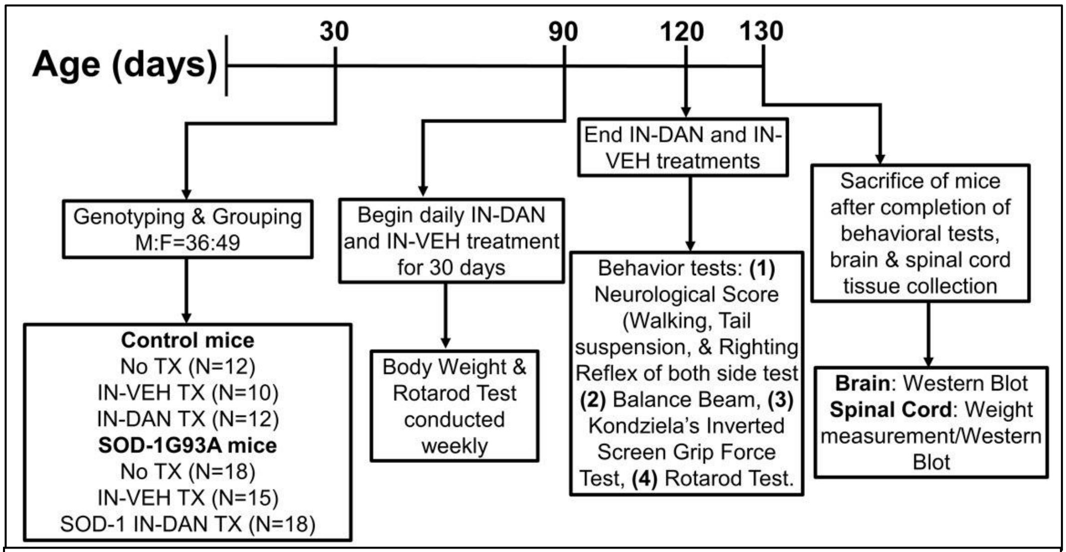


**Supplemental Figure 1. Experimental Design.** No TX: No treatment. IN-DAN: Intranasal dantrolene nanoparticle treatment. IN-VEH: Intranasal vehicle control.

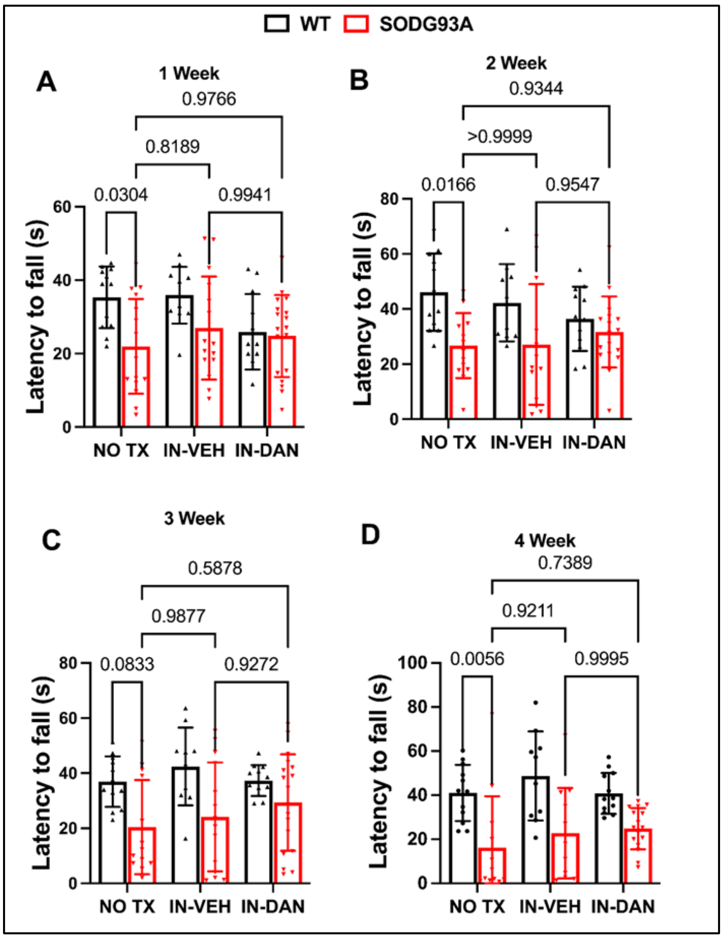


**Supplemental Figure 2. Effects of intranasal dantrolene nanoparticles on motor coordination and balance.** Motor coordination and balance were evaluated weekly over a four-week treatment period using the rotarod test, following the initiation of intranasal dantrolene particle treatment (IN-DAN), vehicle control (IN-VEH), or no treatment (NO TX) between the ages of 90 and 118 days. A longer latency to fall from the rotarod indicates improved motor coordination and balance. At treatment initiation, N = 12 (NO TX), 10 (IN-VEH) and 12 (IN-DAN) for the non-transgenic control mice. N = 18 (No TX), 15 (IN-VEH) and 18 (IN-DAN) for the SOD1-G93A transgenic ALS mice. Data are presented as means ± 95%CI and were analyzed using two-way ANOVA, followed by Tukey's multiple comparison test (MCT).

**Supplemental Video 1 Website: Beam Balance Test**

https://drive.google.com/file/d/14iJB82VOsxZOjKOhDzqY532YrL96PPXM/view?usp=sharing

**Supplemental Video 2 Website:  Grip Strength Test on Inverted Screen**

https://drive.google.com/file/d/1frY2tBQCcZUODTSGzwTnT_edRs_kTwCb/view?usp=sharing

**Supplemental Video 3 Website. Activity Observation**

https://drive.google.com/file/d/100QVvRzTwBDF5XFyfunhtSXp7LVaEbC6/view?usp=sharing
